# Zinc ions prevent α-synuclein aggregation by enhancing chaperone function of human serum albumin

**DOI:** 10.1101/2022.07.04.498643

**Authors:** Samah Al-Harthi, Vladlena Kharchenko, Papita Mandal, Spyridon Gourdoupis, Lukasz Jaremko

**Author notes:** CORRESPONDING AUTHOR Lukasz Jaremko.

## Abstract

Metal ions present in cellular microenvironment have been implicated as drivers of aggregation of amyloid forming proteins. Zinc (Zn^2+^) ions have been reported to directly interact with α-synuclein (AS), a causative agent of Parkinson’s disease and other neurodegenerative diseases, and promote its aggregation. AS is a small intrinsically disordered protein (IDP) i.e., understanding molecular factors that drive its misfolding and aggregation has been challenging since methods used routinely to study protein structure are not effective for IDPs. Here, we report the atomic details of Zn^2+^ binding to AS at physiological conditions using proton-less NMR techniques that can be applied to highly dynamic systems like IDPs. We also examined how human serum albumin (HSA), the most abundant protein in human blood, binds to AS and whether Zn^2+^ and/or ionic strength affect this. We conclude that Zn^2+^ enhances the anti-aggregation chaperoning role of HSA that relies on protecting the hydrophobic N-terminal and NAC regions of AS, rather than polar negatively charged C-terminus. This suggested a previously undocumented role of Zn^2+^ in HSA function and AS aggregation.

## INTRODUCTION

Neurodegenerative conditions are a diverse group of currently untreatable diseases that affect tens of millions of people worldwide (1). The symptoms are characterized by gradual decline of cognitive and locomotor capabilities that are caused by neuronal damage and progressive loss of neurons. Although molecular events that drive neurodegenerative diseases (NDs) remain under intense debate, protein misfolding and aggregation have been proposed as the key molecular drivers of neurodegeneration. An example of one such protein is α-synuclein (AS), an abundant neuronal protein found to aggregate into inclusion bodies, called Lewy bodies, that are characteristic for Parkinson’s disease (PD), dementia with Lewy bodies (DLB) and multiple system atrophy (MSA) (2) (3).

AS is a 140 amino-acid, intrinsically disordered protein (IDP) of unclear physiological function. AS is mainly located in the brain, primarily within presynaptic terminals, and interacts with phospholipids (4) and a range of protein partners (5) (6) (7). Functionally, AS appears to be involved in neurotransmitter release and vesicle trafficking (8); however, whether functionally competent AS is mostly folded (9) (10) (11) (12) or disordered (13) (14) (15) remains unclear. Furthermore, the exact nature of cellular conditions that trigger AS misfolding and aggregation into neurotoxic species and factors that counteract them are also not well understood.

Among many components of the cellular microenvironment, metal ions have emerged as potentially critical regulators of AS structure and aggregation. One of the most prominent examples in this context are zinc ions (Zn^2+^) that have been shown to accelerate AS amyloid fibril formation (16). Based on a wide range of biophysical techniques (UV-VIS CD spectroscopy (17), fluorescence spectroscopy (18), and mass spectroscopy (19)) and in vivo (20) studies, Zn^2+^ was proposed to bind to both the C-terminal and N-terminal domains of AS with low millimolar affinity. However, the results of these biophysical studies have recently been called into question, given that physiological Zn^2+^ concentrations in synapses (10-50 μM) are significantly lower than the mM concentrations used in in vitro studies (21) (22) (23) (24) (25). Additionally, AS purified from natural sources (human red blood cells and human brain tissue samples), showed no presence of Zn^2+^ (26). Therefore, despite extensive research, the role of Zn^2+^ in AS physiology and pathology is yet to be established.

In order to examine this question further, we focused on the study of how Zn^2+^ ions affect AS using specially designed nuclear magnetic resonance (NMR) spectroscopy pulse sequences that enable monitoring IDPs under physiologically relevant conditions. In general, NMR spectra of IDPs suffer from poor resolution and extensive resonance overlaps due to unstructured and conformationally dynamic nature of IDPs. To address this problem, we employed ^13^C-detected 2D NMR experiments (CBCACO, CACO, CON), which have been shown as effective in yielding structural information in the recent study of the interaction between AS and Ca^2+^ (27). More specifically, we focused on providing further insights into how Zn^2+^ ions affect extracellular AS, given that this form of AS was proposed to be essential for neuron-to-neuron transmission and disease progression (28) In this context, we examined the three-component system of AS, Zn^2+^, and human serum albumin (HSA). HSA is the most abundant protein in the blood (60%) (29) (30), and due to its capacity for binding several different ions such as Zn^2+^, and other molecules, including many proteins (31) (32) (33), it is considered to be one of the most important molecular transporters in biology (31) (34) (35) (36). In addition to the role as a transporter, recent analysis revealed that HSA also serves as an aggregation inhibitor with a chaperon-like activity for proteins that tend to oligomerize like the amyloidogenic peptide Aβ (37) (38) (39) (40) (41) (42) (43) and AS (15) (44) (45). Our analysis of the three-component system of AS, HSA and Zn^2+^, under physiological, and no-salt conditions revealed Zn^2+^-dependent chaperoning effects of HSA on AS fibril formation and inhibition of AS aggregation. Thus, our study documents an unexpected role of Zn^2+^ as an enhancer of HSA chaperone activity.

## RESULTS

### Effect of Zn^2+^ and/or human serum albumin on AS fibrillization kinetics under different ionic strength conditions

We used the well-established ThT fluorescence assay **(Figure 1A)** to monitor the effect of Zn^2+^ concentration and/or defatted HSA (DE-HSA). We chose to initially focus on DE-HSA because AS itself has affinity for fatty acid binding; therefore, we wanted to study AS/HSA and AS/Zn^2+^/HSA binding separately from any potential effects of fatty acids. We measured how increasing concentration of Zn^2+^ affect fibrillization kinetics of AS under physiological conditions (37 °C, 20 mM HEPES, pH 7.4, 140 mM NaCl), as well as no salt conditions (37 °C, 20 mM HEPES, pH 7.4, 0 mM NaCl) **(Figure 1B)**. Under both sets of conditions, increasing Zn^2+^ concentration led to increase in AS fibrillization. We also examined how addition of DE-HSA or Zn^2+^/DE-HSA affects AS fibrillization kinetics and observed dramatic decrease in fibrillization under physiological conditions when either DE-HSA alone or Zn^2+^/DE-HAS was added **(Figure 1C)**. Although addition of increasing amounts of Zn^2+^ led to accelerated fibrillization, addition of DE-HSA into AS containing buffer did not initiate any AS oligomerization process, and the fibrillization remained at halt even after addition of increasing amounts of Zn^2+^. We repeated the same procedure under low ionic strength conditions (37 °C, 20 mM HEPES, pH 7.4, 0 mM NaCl) **(Figure 1D)**. The fibrillization of AS reached the plateau faster, at a lower concentration of Zn^2+^. Furthermore, the fibrillization process was not completely halted in the presence of increased amounts of DE-HSA and increased amounts of DE-HSA and Zn^2+^.

**Figure 1.**
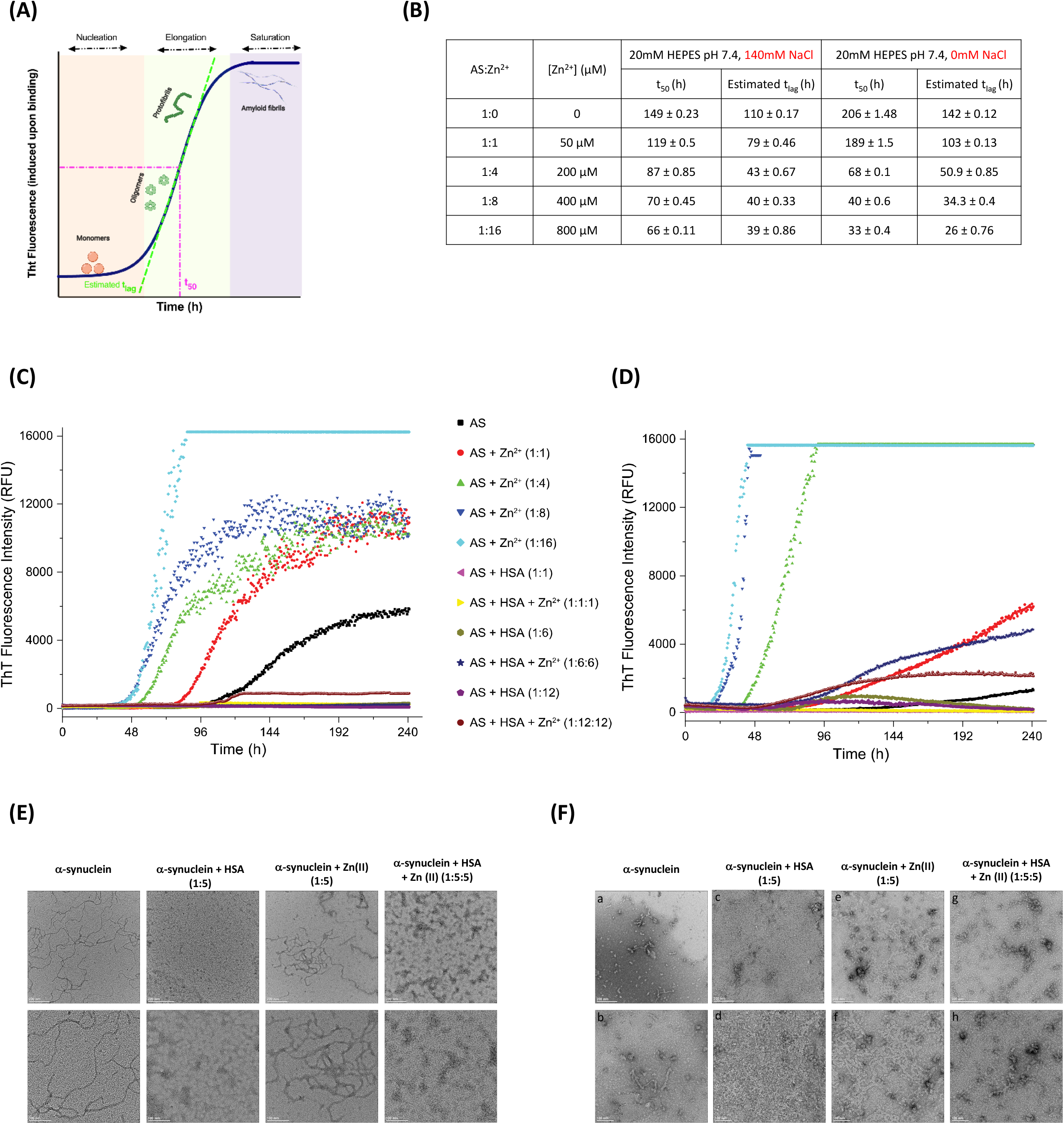
Fibrillization kinetics of AS under pH 7.4, low and high ionic strength in the presence and absence of Zn^2+^ and/or defatted human serum albumin (DE-HSA). (A) The nucleation dependent polymerization reaction of amyloid fibril formation causes a sigmoidal growth curve with nucleation, elongation, and saturation phases. (B) Table highlighting the estimated t_50_ and t_lag_ parameters for two experimental conditions setup. (C) AS fibrillization in the presence of increasing amount of Zn^2+^ (AS:Zn^2+^, 1:1, 1:4, 1:8, 1:16 molar ratio) and increasing amount of HSA and/or Zn^2+^ under high-ionic strength (140 mM NaCl) and (D) under low-ionic strength (0 mM NaCl). (E) TEM images of fibrils formation of (a, b) AS (WT); (c, d) AS in the presence of HSA (1:5); (e, f)) AS in the presence of Zn^2+^ (1:5); (g, h)) AS in the presence of HSA and Zn^2+^ (1:5) under high-ionic strength and (F) under low-ionic (Scale bar: upper raw 200nm and lower raw 100nm).

To examine the effect of fatty acids on AS aggregation, we repeated the ThT fluorescence assay with FA-HSA, where FA represents either caproic or palmitic acid. We observed that addition of FAs did not change fibrillization kinetics (Supplementary Material Figure S1).

We observed that ionic strength affected fibrillization kinetics, which led us to examine whether the kinetic effect translate into distinct morphological features. We performed transmission electron microscopy (TEM) imaging of AS alone, AS with DE-HSA, AS with Zn^2+^, and with combination of DE-HSA and Zn^2+^ in the presence **(Figure 1E)** and absence **(Figure 1F)** of 140 mM of NaCl. The results were in agreement with the ThT fluorescence assay, as we observed that fibrillization of AS was enhanced in the presence of Zn^2+^, resulting in larger and more condensed fibrils at both high and low ionic strength conditions compared to AS alone. DE-HSA inhibited the formation of fibrils under all conditions (with or without Zn^2+^, in the presence and absence of 140 mM NaCl); however, as noted above, inhibitory effects were more pronounced under physiological (high ionic strength) conditions. Morphologically, we noted that fibrils formed under physiological ionic strength conditions are longer **(Figure 1E)**, while the fibrils formed at the low ionic strength conditions are thicker and more condensed **(Figure 1F)**. Same results were consistently observed in several replications of all conditions (Supplementary Material Figures S4 and S5).

Taken together, this data illustrates that AS fibrillization kinetics is sensitive to Zn^2+^ concentrations, and greatly accelerated when Zn^2+^ ions were added alone. However, addition of DE-HSA and FA-HSA slowed down the process significantly. We also observed that this process was sensitive to ionic strength, as these inhibitory effects were somewhat diminished under no-salt conditions. These differences in kinetics translated into distinct morphology of fibrils formed in high-salt versus no-salt conditions, suggesting that electrostatics play key roles in guiding the process of AS fibril formation.

### Proton-less NMR maps Zn^2+^ interaction with AS under physiological conditions

We employed proton-less NMR experiments to map AS residues affected by Zn^2+^ binding at body temperature of 37 °C. From the analysis of 2D CON **(Figure 2A)** and 2D CBCACO **(Figure 2B)** NMR spectra of the titration of Zn^2+^ to AS, we identified the residues experiencing chemical shift change in the fast exchange regime upon addition of increasing amounts of Zn^2+^. We observed a subset of residues that displayed clear chemical shift changes, indicative of specific interaction. To get a more quantitative view of the changes, we calculated chemical shift perturbation (CSP) on per residue basis based on the 2D CON NMR monitored titration. The CSP results highlight involvement of both N-and C-terminal residues in mediating Zn^2+^ interactions **(Figure 2C)**. To get a more molecular view of the Zn^2+^ binding, we mapped the residues with the largest CSP onto the structures of micelle-bound AS (*1XQ8*.*pdb*) and AS fibrils (*2N0A*.*pdb*) **(Figure 2D)**. We observed that Zn^2+^ anchoring regions (indicated in orange, **Figure 2D)**, map onto both structured and disordered sections of the structure. Collectively, our data support existence of well-defined and specific Zn^2+^ binding sites on AS represented primarily by aspartates and glutamates located in N-and C-terminal regions, but not within non-amyloid b component (NAC) **(Figure 2C)**. This suggests that specific Zn^2+^ ion binding accelerates AS fibrillization by reducing the enthalpic penalty of fibrillization by masking the negative charges and, hence, stabilizing oligomerization intermediates and/or fibrils.

**Figure 2.**
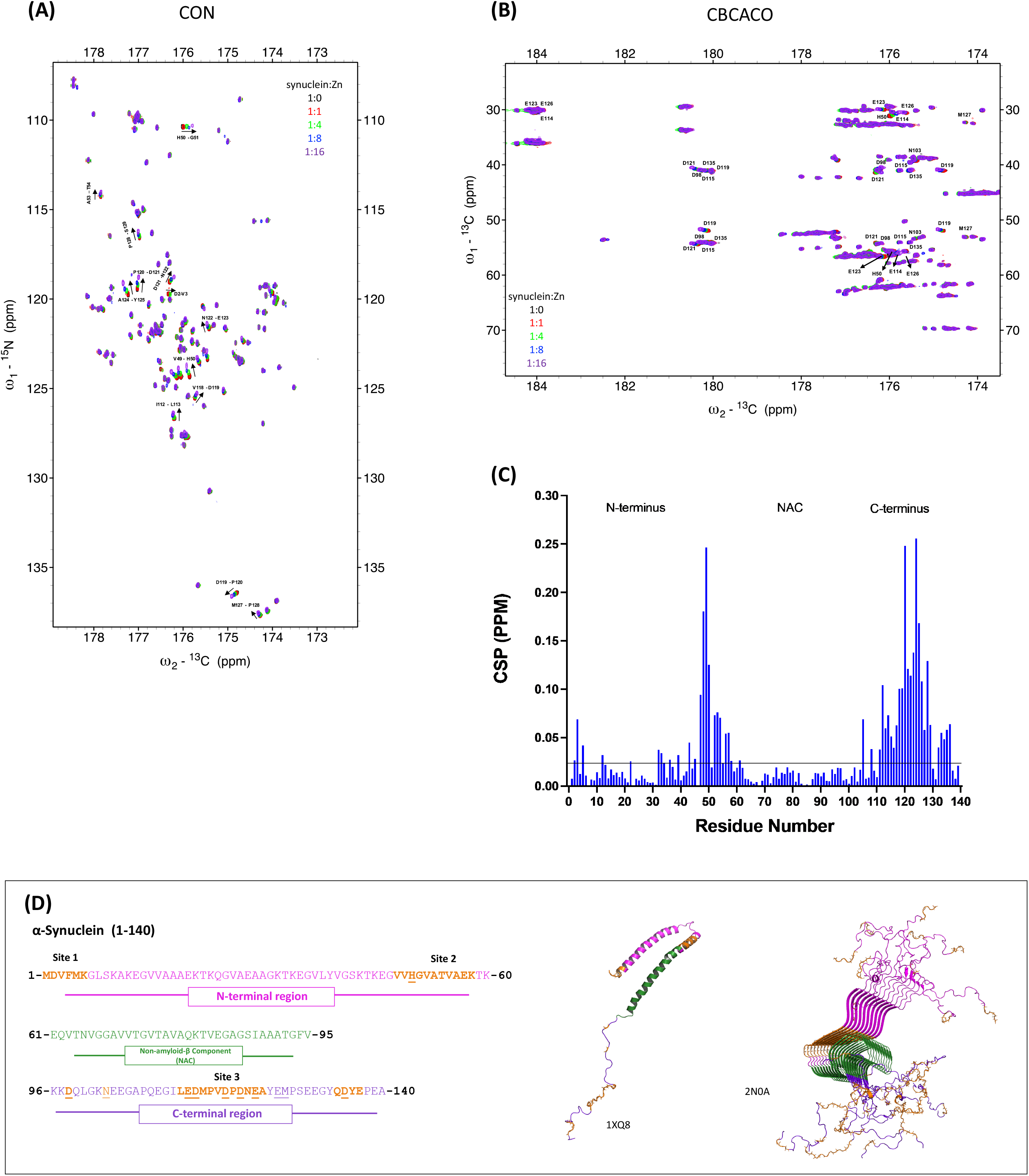
The atomic level proton-less NMR-monitored Zn^+2^ binding to AS under physiological conditions. (A) Overlaid 2D CON spectrum and (B) 2D CBCACO spectrum of AS (150μM) in the absence (black) and presence of 150 μM (red), 600 μM (green), 1200 μM (blue) and 2400 μM Zn^+2^ (purple). (C) Chemical shift perturbation of CON signals upon addition of Zn^2+^. (D) Primary sequences of AS (1-140) highlighting the N-terminal, NAC and C-terminal domains and PDB structures of Human micelle-bound AS (1XQ8) and AS fibrils (2N0A). The highlighted residues represent the main Zn^+2^ anchoring sites reported upon the interactions with Zn^+2^ ions (highlighted in orange obtained from CON spectrum and underlined residues from CBCACO spectrum).

### Zn^2+^ ions enhance HSA hydrophobic binding to AS at physiological salt conditions

To examine how HSA affects AS, we conducted 2D CON NMR titration experiments for AS with DE-HSA **(Figure 3A)** and with DE-HSA+Zn^2+^ **(Figure 3B)** under physiological conditions (37 °C, 20 mM HEPES, pH 7.4, 140 mM NaCl). We plotted the intensity ratios (I/I_0_) of the 1-140 non-overlapping cross peaks which we identified in the 2D CON NMR titration spectra and we validated that HSA binds to AS by observing the intensity profile regions with lower I/I_0_ ratios indicating higher levels of perturbation **(Figure 3C, D)**. We noticed that DE-HSA alone interacts with the initial part of N-terminal domain but not with NAC or C-terminal domain **(Figure 3C)**. Interestingly, Zn^2+^ binding to DE-HSA greatly enhances interactions with AS, especially with N-terminal and NAC domains, as indicated by much lower I/I_0_ ratios **(Figure 3D)**. Given that both N-terminal and NAC domains are primarily hydrophobic and that high ionic strength diminishes the electrostatic effects, our binding data suggests that Zn^2+^ ion binding promotes hydrophobic interactions. A gradient representation of I/I_0_ of the affected sites is shown on the PDB structures of micelle-bound AS and AS fibrils, illustrating that the perturbations are localized, suggesting the specific non-random nature of these interactions **(Figure 3E, F)**. These results were further confirmed by 2D CBCACO NMR spectra (Supplementary Material Figure S2). Taken together, these studies indicate that HSA inhibits the AS aggregation by both affecting AS directly, and by reducing the access of Zn^2+^ to AS thus preventing oligomerization of monomeric AS.

**Figure 3.**
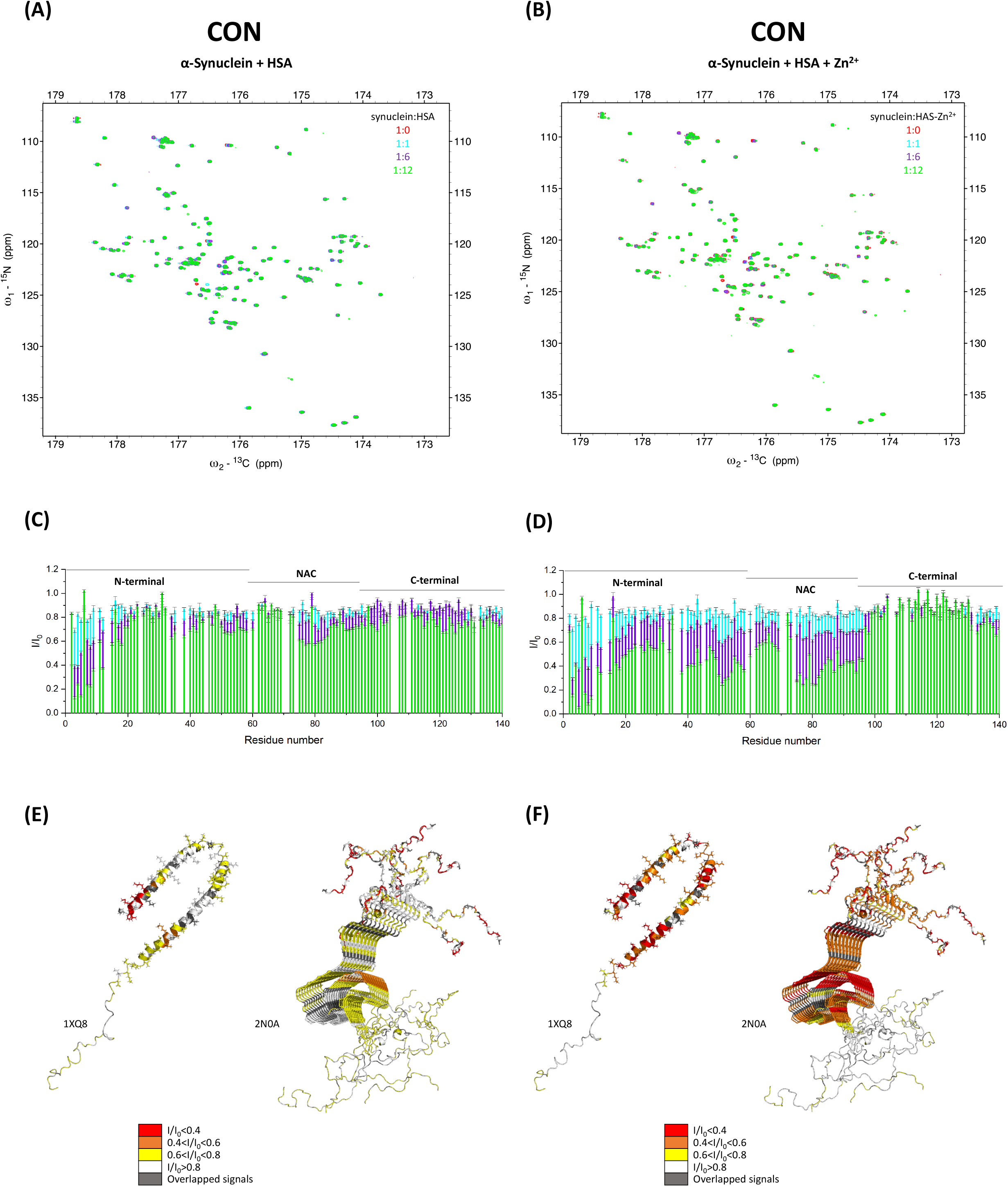
Impact of Zn^+2^ on AS binding to HSA under high ionic strength. (A) Overlaid 2D CON spectrum of AS (150 μM) in the absence (red) and presence of 150 μM (cyan), 900 μM (purple), and 1800 μM (green) HSA and (B) HSA-Zn^+2^. (C) The I/I_0_ ratios of 1-140 non-overlapping cross peaks as a function of protein sequence at the presence of HSA and (D) HSA-Zn^+2^. (E) PDB structures of Human micelle-bound AS (1XQ8) and AS fibrils (2N0A) highlighting the AS residues that are affected by addition of HSA and (F) HSA-Zn^+2^ at the presence of 140 mM NaCl. Largely affected (red), significantly affected (orange), mildly affected (yellow), not affected (white) and overlapped signals (gray).

### Lower ionic strength favors electrostatic interactions between HSA and AS

To investigate the effect of salt concentration we repeated the same analysis (2D CON) under low ionic strength conditions (37 °C, 20 mM HEPES, pH 7.4, 0 mM NaCl) **(Figure 4A, 4B)**. In line with the ThT fluorescence assay and the TEM images, the intensity profiles of AS titrated with DE-HSA **(Figure 4C)** and with Zn^2+^-bound DE-HSA **(Figure 4D)** validated that the hydrophobic binding of both DE-HSA alone and DE-HSA bound with Zn^2+^ on AS is lower than that at physiological ionic strength conditions. Under low ionic strength conditions, we observed that addition of DE-HSA induces perturbations only at the C-terminal domain of AS **(Figure 4C)**. Addition of Zn^+2^ diminished these effects, while resulting in more global distribution of smaller I/I_0_ ratio changes throughout the AS **(Figure 4D)**. We mapped I/I_0_ ratios onto micelle-bound AS and AS fibrils’ structure, and observed that the impact is much smaller and concentrated to the flexible C-terminal region **(Figure 4E)**. We also recorded the 2D CBCACO NMR spectra of the same titration combinations, which are in agreement with 2D CON data (Supplementary Material Figure S3). The aforementioned are also verified by comparing the ITC profiles of AS with DE-HSA versus Zn^2+^ and DE-HSA alone versus Zn^2+^ at physiological high ionic strength conditions (Supplementary Material Figure S6). Therefore, we conclude that under low ionic strength conditions binding events among different entities in the solution are driven by electrostatics, and therefore unlikely to be specific or reflective of physiologically relevant binding events.

**Figure 4.**
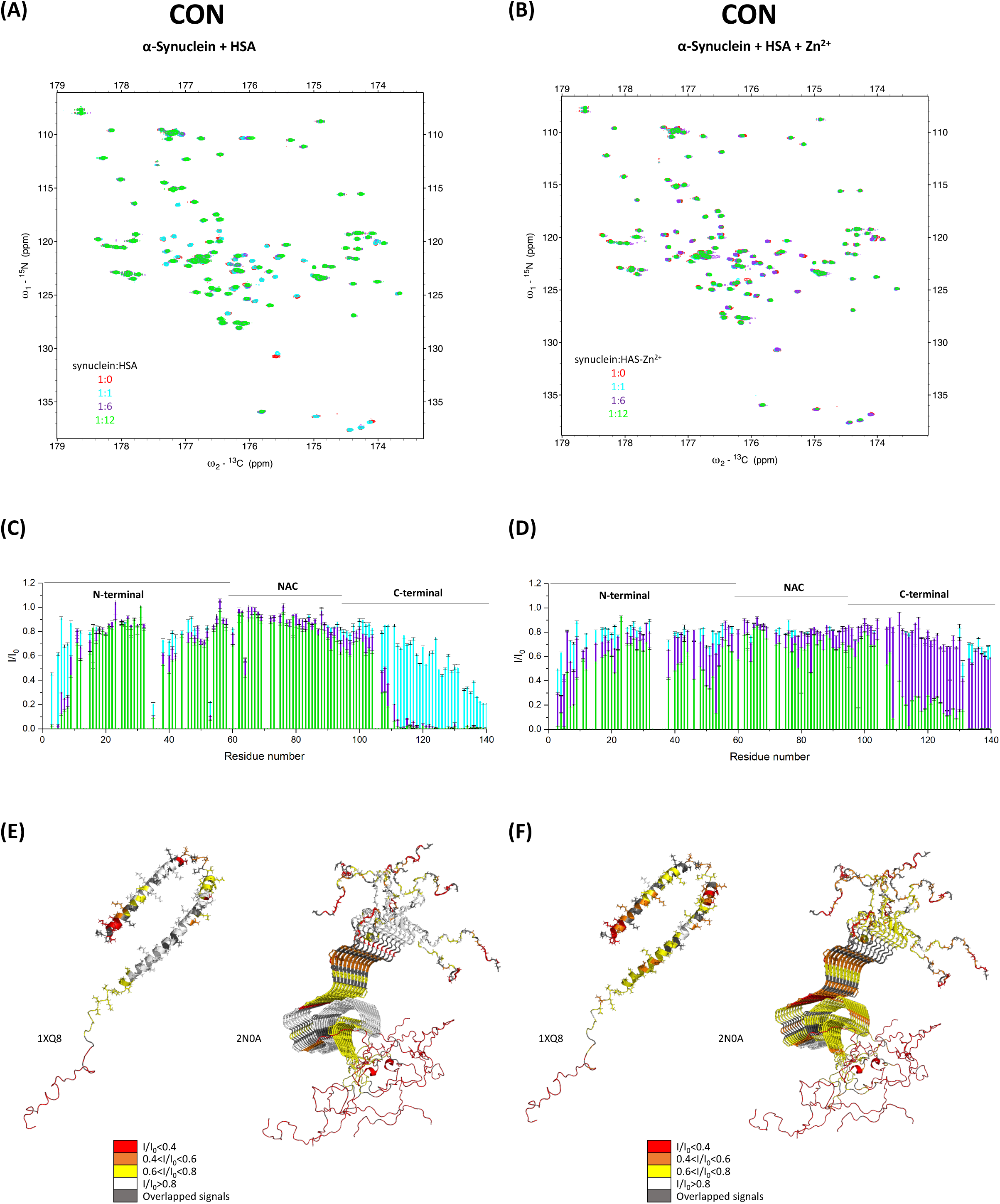
Impact of Zn^+2^ on AS binding to HSA under low ionic strength. (A) Overlaid 2D CON spectrum of AS (150 μM) in the absence (red) and presence of 150 μM (cyan), 900 μM (purple), and 1800 μM (green) HSA and (B) HSA-Zn^+2^. (C) The I/I_0_ ratios of 1-140 non-overlapping cross peaks as a function of protein sequence at the presence of HSA and (D) HSA-Zn^+2^. (E) PDB structures of Human micelle-bound AS (1XQ8) and AS fibrils (2N0A) highlighting the AS residues that are affected by addition of HSA and (F) HSA-Zn^+2^ at the presence of 0 mM NaCl. Largely affected (red), significantly affected (orange), mildly affected (yellow), not affected (white) and overlapped signals (gray).

## DISCUSSION

Taken together, our study investigated the interaction of AS with Zn^2+^. Zinc ions have been implicated as important modulators of fibrillization of amyloid forming proteins (19), and yet exact mechanisms of how these ions may be driving fibrillization remain incompletely understood. Furthermore, given that neuron-to-neuron transmission seems to depend on the presence of extracellular AS, we examined fibrillization properties of AS in the presence of HSA, the most abundant protein in human blood that has been proposed to act as an AS chaperone and prevent its aggregation (44). Our fibrillization kinetics measurements showed that AS oligomerization is dramatically accelerated by increasing concentrations of Zn^2+^ ions; however, under physiological ionic strength, addition of HSA/ Zn^2+^ had an opposite effect, slowing down AS aggregation. This suggests an intriguing possibility that Zn^2+^ enhances chaperoning capabilities of HSA.

To provide a more granular insight into these effects, we used proton-less NMR methods (2D CON and 2D CBCACO) that allowed us to record per residue-based data on AS, despite its IDP nature. Through these NMR titration experiments we learned that both Zn^2+^ and HSA bind to AS by engaging specific, well-defined regions through mostly hydrophobic interactions. Importantly, addition of Zn^2+^ enhances HSA/AS binding effects, further supporting our proposed hypothesis that Zn^2+^ strengthens HSA chaperoning capabilities under physiological ionic strength conditions. More specifically, based on the CSP data we observed that Zn^2+^ binds AS via three well-defined sites, mostly by coordinating negatively charged aspartate and glutamate residues. This suggests that Zn^2+^ binding neutralizes some of the charged regions of the AS, which may drive accelerated aggregation by decreasing some of the electrostatic repulsion between individual AS molecules. This notion is further supported by ionic strength effects, whereby we observed that under low ionic strength conditions the effects of Zn^2+^ binding on fibrillization kinetics are even more pronounced.

We also observed that ionic strength affects HSA and AS binding. Under physiological conditions, HSA interacts mostly with N-terminal and NAC regions of AS. Given the mostly hydrophobic nature of these regions, our results suggest that this binding is mediated through hydrophobic interactions. Addition of Zn^2+^ seems to mask charges and enhance hydrophobicity. To explore the impact of hydrophobicity and charge surface further, we modeled these parameters onto human micelle-bound AS, HSA (*1AO6*.*pdb*) and Zn^2+^-bound HSA (*5IJF*.*pdb*) **(Figure 5)**. We noted that the changes at the charge of the surface of HSA upon Zn^2+^ binding do not translate to proportional changes on the hydrophobicity of HSA. Thus, the exact details of the mechanism remain to be determined. Finally, our results strongly argue for the need to conduct AS aggregation and binding experiments under conditions that closely mirror human physiological conditions to prevent potential misleading findings of non-biological relevance.

**Figure 5.**
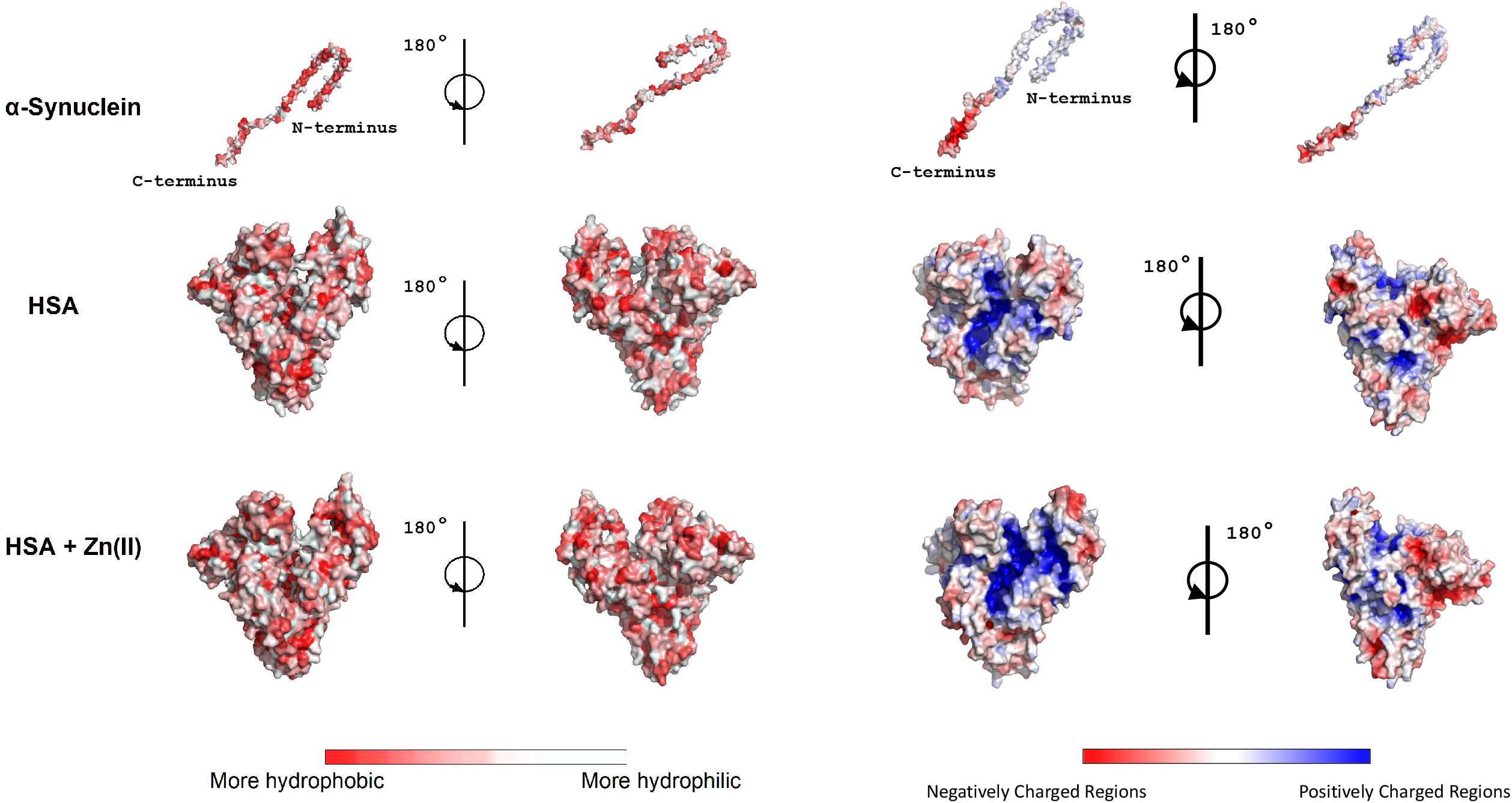
The interplay of hydrophobic and electrostatic interactions of AS, HAS and Zn^2+^ interactions. Surface hydrophobicity calculations (first panel) and surface charge representation (second panel) of human micelle-bound AS (PDB: 1XQ8), HSA (PDB: 1AO6) and Zn^2+^-bound HSA (PDB: 5IJF). The electrostatic surfaces were generated with the APBS plugin of PyMOL2.

## MATERIALS AND METHODS

Materials. De-fatted Human Serum Albumin (DE-HSA) lyophilized powder was purchased from Sigma-Aldrich (product no. A3782-1G; >99% pure base on gel electrophoresis). Less than 0.02% fatty acids are present, globulin free and has been tested negative for HIV I, HIV II, HCV, and HBsAg. HSA was dissolved in 20 mM HEPES pH 7.4, 140 mM NaCl to obtain the required concentration for each experiment. The exact HSA concentration was determined by UV absorbance using NanoDrop™ One Spectrophotometer using molar extinction coefficient of 35,700 M^-1^cm^-1^ at 280 nm. The protein was then dialyzed overnight at 4 °C against respective buffer in order to remove impurities. Palmitic and caproic acid were also purchased from Sigma-Aldrich (product no. P0500-10G and 153745-100G respectively).

Fatty acid/HSA Complexes Preparation. Fatty acids (FAs) were dissolved in 50% ethanol and heat up to 70 °C to obtain clear stock solutions with a concentration of 10 mM prior to each experiment. Solutions were diluted by Tris-NaCl buffer to obtain 1 mM concentration and warmed up to about 50 °C in order to facilitate fatty acid dispersion. Afterwards, the solutions were allowed to evaporate at room temperature and slightly cool down, and degassed for 20 minutes to eliminate the ethanol before the addition to HSA. FA-HSA complexes were prepared by the addition of the appropriate volumes of FAs stock to constant protein volume to obtain the desired FA:HSA molar ratios of 1:1, 2:1, 4:1 and 8:1. The mix incubated at 500 RPM shaker for 2 hours at 30 °C to enable fatty acids binding to HSA. Monitoring by 1D ^1^H NMR we confirmed that the presence of ethanol in the FAs was less than 1% compared to the 5% ethanol initially in the sample i.e., the processing of the sample achieved maximum ethanol evaporation.

Recombinant AS expression and purification. Double-labeled (^13^C, ^15^N) and unlabeled AS were used in this study. Untagged AS was expressed in BL21-DE3 RIL E.coli cells grown in either ^15^NH_4_Cl, ^13^C-glucose containing M9 minimal media or in 2X YT media. Bacterial cultures growing at 37 °C with shaking were induced with 1 mM IPTG at OD_600_ 0.5-0.6. After continued growth at 37 °C for 4 hours, cells were harvested and lysed in 20 mM Tris, pH 8, 25 mM NaCl, 1 mM EDTA containing buffer. The lysate was boiled at 80 °C for 15 minutes and allowed to cool down on ice followed by centrifugation at 15,000 g at 4 °C for 30 minutes. The clarified lysate was loaded onto HiTrap Q HP column pre-equilibrated in 20 mM Tris, pH 8, 25 mM NaCl, 1 mM EDTA buffer. AS was eluted from the Q column using a linear gradient of 25 mM - 1 M NaCl buffered with 20 mM Tris, pH 8 over 5 column volumes. The AS containing fractions were pooled and dialyzed against 20 mM sodium phosphate, pH 6.5, 100 mM NaCl, 5 mM EDTA buffer and subjected to size exclusion chromatography using Superdex 75 10/300 column pre-equilibrated in the same buffer. AS containing fractions were pooled and stored at -20 °C until further use.

ThT Aggregation Assay. AS fibrillization in the presence and/or absence of Zn^2+^ and HSA were examined using ThT assay. First, AS fibrillization was examined under 20 mM HEPES, pH 7.4, 140 mM NaCl condition setup. 50 μM ThT dye and 50 μM wild type AS monomers were mixed with increasing amount of ZnCl_2_ (AS:ZnCl_2_: 1:0, 1:1, 1:4, 1:8, 1:16). Then 50 μM ThT dye and 50 μM wild type AS monomers were mixed with DE-HSA, DE-HSA in complex with Caproic acid and DE-HSA in complex with Palmitic acid in the presence and absence of increasing amount of ZnCl_2_. AS fibrillization was also examined under 20 mM HEPES, pH 7.4, 0 mM NaCl condition setup. 50 μM ThT dye and 50 μM wild type AS monomers were mixed with increasing amount of ZnCl_2_ (AS:ZnCl_2_: 1:0, 1:1, 1:4, 1:8, 1:16). Then 50 μM ThT dye and 50 μM wild type AS monomers were mixed with DE-HSA in the presence and absence of increasing amount of ZnCl_2_. Each sample was split into triplicate of 200 μL and placed in the wells of a sealed, flat clear-bottomed 96-well plate (Corning). ∼ 15 Glass beads were added to each well (1 mm glass bead, Sigma-Aldrich) in order to improve mixing and enhance homogeneity among replicates. The plate we inserted into SpectraMax M5 Microplate Reader and were incubated at 37 °C, for 29 min and subjected to 1 min shaking. The ThT fluorescence was read by excitation wavelength at 440 nm and emission wavelength at 485 nm for over 240 hours with 30 min intervals. The three replicates of each sample measured were averaged, corrected and analyzed.

The data were fitted to the Boltzmann equation (a sigmoidal function) as given by:

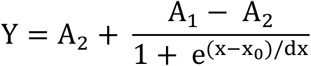

A_1_ is the minimum signal, A_2_ is the maximum signal, x_0_ is the time at which the change in signal is 50% (t_50_), dx is the time constant, and the lag time is estimated by:

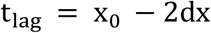

TEM imaging. AS fibrillization alone, and in the presence of AS:HSA (1:5 ratio), AS:Zn^2+^ (1:5 ratio) and AS:HSA:Zn^2+^ (1:5:5 ratio) were examined using high-resolution TEM imaging. AS fibrillization was examined under 20 mM HEPES, pH 7.4, 140 mM NaCl and 20 mM HEPES, pH 7.4, 0 mM NaCl conditions setups. The prepared samples for both conditions were incubated for 10 days at 37 °C without shaking. Then, 10 μL of each sample was pipetted onto glow-discharged carbon coated copper grid and incubated for 1 min. The excess sample was drawn off using filter paper followed by washing twice with dH_2_O. Then 2% uranyl acetate was used to stain the samples for 30 seconds followed by air drying the grids. Imaging was performed on Titan CT TEM at KAUST Core Labs.

Isothermal Titration Calorimetry (ITC). ITC experiments were carried out with MicroCal PEAQ (Malvern Instruments, MA) at 25 °C at 20 mM HEPES, pH 7.4, 140mM NaCl and at 20mM HEPES, pH 7.4, 0mM NaCl conditions setups. This buffer was chosen mainly due to its negligible interaction with Zn^2+^. All titrated proteins, i.e. AS (monomer), HSA, and AS:HSA mixture were kept in the cell with maintained concentration of 50 μM while ZnCl_2_ was kept in the syringe at 1mM concentration. Measurements with 25 injections of initial 0.2 μL injection volume with an initial delay of 60 seconds followed by 2 μL over 20 seconds duration and spacing of 180 seconds were performed. The binding curves were fitted assuming a single-site binding model using the analysis software provided by the manufacturer.

NMR spectroscopy. NMR data for sequence specific resonance assignment of AS were acquired at 700 and 800 MHz Bruker Avance NEO spectrometers equipped with cryoprobe, running Topspin version 4.0.7. The measurements were acquired at 288 K, 570 μM of ^13^C^15^N-labeled AS in 20 mM phosphate buffer (NaPi), pH 6.5, 100 mM NaCl and 1% D_2_O for the lock signal. Backbone ^1^H, ^13^C, ^15^N resonances were assigned by recording and analyzing TROSY-based triple resonance three-dimensional (3D) experiments HNCA, HNCO, HNCACO, HNCACB, and CBCACONH. NMR data processing, analysis and assignment of the data were performed using Topspin ver. 4.0.7 and CARA (http://cara.nmr.ch/) software.

NMR Monitored Titration Experiments. For the titration experiments, 150 μM of ^13^C^15^N-labeled AS in 20mM HEPES pH, 7.4, 140mM NaCl and 1% D_2_O titrated with increasing amount of ZnCl_2_ (AS:ZnCl_2_: 1:0, 1:1, 1:4, 1:8, 1:16). ZnCl_2_ solution was prepared with a final concentration of 100 mM in the same buffer. The second set of samples were prepared by the addition of HSA and/or ZnCl_2_ to 150 μM of ^13^C^15^N-labeled AS in 20mM HEPES, pH 7.4, 140 mM NaCl. The AS:HSA and AS:HSA:Zn^2+^ ratios were 1:1 and 1:1:1, 1:6 and 1:6:6, 1:12 and 1:12:12. The second set of samples with the same ratios were also prepared in 20 mM HEPES, pH 7.4, 0 mM NaCl. Measurements of all the samples were carried out at 37 °C on a 700 MHz Bruker Avance NEO spectrometer equipped with cryoprobe, running Topspin version 4.0.7. ^1^H-^15^N TROSY-HSQC, HCACO, HCBCACO, and HCACON spectra were collected for each sample. NMR data processing and interpretation were performed using Topspin ver. 4.0.7, SPARKY (https://www.cgl.ucsf.edu/home/sparky/) and CARA (http://cara.nmr.ch/) software.

## ACKNOWLEDGMENTS

This publication is based upon work supported by the King Abdullah University of Science and Technology (KAUST) Office of Sponsored Research (OSR) under Award No. OSR-CRG2018-3792 and Award No. OSR-CRG2019-4088 (L.J.).

## AUTHOR CONTRIBUTIONS

The project was conceived and supervised by L.J. and S.Al-H. Experimental studies and analysis were done by S.Al-H., V.K and S.G. The double labeled α-synuclein sample for NMR was prepared by P.M. The manuscript was written by S.Al-H, LJ, S.G., and through contributions of all other coauthors. All authors have given approval to the final version of the manuscript.

## CONFLICT OF INTEREST

The authors declare no conflict of interest.

## FUNDING SOURCES

Award No. OSR-CRG2018-3792

Award No. OSR-CRG2019-4088

## SUPPLEMENTARY FIGURES

**Figure S1 -** AS fibrillization in the presence of DE-HSA and FA-HSA (HSA in complex with Caproic acid and Palmitic acid) and Zn^2+^ to examine the impact of FAs on AS aggregation.

**Figure S2 -** NMR analysis of HSA and/or Zn^2+^ titration to AS under 20 mM HEPES pH 7.4, 140 mM NaCl. Top spectrum overlaid 2D CBCACO spectrum of AS (150 μM) in the absence (red) and presence of 150 μM (cyan), 900 μM (purple), and 1800 μM (green) HSA. Bottom spectrum overlaid 2D CBCACO spectrum of AS (150 μM) in the absence (red) and presence of 150 μM (cyan), 900 μM (purple), and 1800 μM (green) HSA and Zn^2+^.

**Figure S3 -** NMR analysis of HSA and/or Zn^2+^ titration to AS under 20 mM HEPES pH 7.4, 0 mM NaCl. Top spectrum overlaid 2D CBCACO spectrum of AS (150 μM) in the absence (red) and presence of 150 μM (cyan), 900 μM (purple), and 1800 μM (green) HSA. Bottom spectrum overlaid 2D CBCACO spectrum of AS (150 μM) in the absence (red) and presence of 150 μM (cyan), 900 μM (purple), and 1800 μM (green) HSA and Zn^2+^.

**Figure S4 -** TEM images of fibrils formation of AS in the presence of HSA (1:5), Zn^2+^ (1:5) and Zn^2+^-HSA (1:5) under high-ionic strength (Scale bar: upper raw 200 nm and lower raw 100 nm).

**Figure S5 -** TEM images of fibrils formation of AS in the presence of HSA (1:5), Zn^2+^ (1:5), and Zn^2+^-HSA (1:5) under low-ionic strength (Scale bar: upper raw 200 nm and lower raw 100 nm).

**Figure S6 -** ITC titration of 2.7 mM ZnCl_2_ into 100 μM AS and HSA (blue) and only HSA (magenta) in 20 mM HEPES pH 7.4, 140 mM NaCl at 25°C.

## ABBREVIATIONS

AS: α-synuclein
HSA: human serum albumin
DE-HSA: defatted human serum albumin
FA-HSA: human serum albumin with fatty acids
ITC: isothermal calorimetry
NMR: nuclear magnetic resonance
BBB: blood-brain barrier
PD: Parkinson’s disease
DLB: dementia with Lewy bodies
MSA: multiple system atrophy
CSF: cerebrospinal fluid
Q-Alb: CSF/serum albumin ratio
ThT: Thioflavin T fluorescence assay
TEM: transmission electron microscopy imaging

